# The Role of 1-*O*-Acylceramide NP in Structural Organization and Permeability of the Stratum Corneum Lipid Matrix

**DOI:** 10.1101/2022.12.06.519381

**Authors:** Moon Young Yang, Eun Ok Lee, Chang Seo Park, Yoon Sung Nam

## Abstract

The extracellular lipid matrix in the stratum corneum (SC) is crucial for generating a skin barrier (permeability) function. The lipid matrix contains three major components; ceramides, cholesterol, and free fatty acids. The broad diversity of ceramides depends on their molecular structures (*e.g*., hydroxylations and chain lengths) and plays a critical role in maintaining the structural integrity of the lipid matrix. Although recent studies identified a new subclass of ceramide, 1-*O*-acylceramide NP (CerENP), its precise role in the lipid matrix of SC is still elusive. Herein, we investigate the role of CerENP on the structure and permeability of the SC by molecular dynamics simulations. Our results suggest that the CerENP molecules induce a denser lipid matrix in the lateral dimension in the long periodicity phase model with a bilayer-slab- bilayer structure. Moreover, ethanol permeability analysis indicates that CerENP can suppress molecular permeability through the lipid matrix. This study provides insight into the role of a new subclass of ceramide in the SC, which can lead to our better understanding of skin organization and disease-related barrier dysfunction.

## 1. INTRODUCTION

The outer layer of the skin, stratum corneum (SC), is a primary barrier against the percutaneous penetration of exterior molecules and water loss across the skin. The SC is mainly composed of corneocytes (differentiated dead keratinocytes) surrounded by a lipid matrix,^1^ which contains three major components, ceramides (Cers), cholesterol (Chol), and free fatty acids (FFA) in an about equimolar ratio,^2^ providing the essential elements of the skin barrier.^3^ Understanding the structural organization of the lipid matrix in the SC is essential to rationalize and control skin permeability as the lipid matrix is continuous throughout the SC and determines the permeation rate of various molecules.^4^

Although the exact structure of the SC is still unknown, there has been extensive efforts to understand the function and structure of the lipids as well as their chemical and biophysical properties. The structure of the lipid matrix in the SC has been investigated using various experimental methods including electron microscopy,^9^ X-ray diffraction,^10^ infrared spectroscopy,^11^ and neutron diffraction.^12^ Also, the ^2^H NMR has been used to quantitatively describe the phase behavior of a deuterated species in a complex lipid mixture.^13,14^ The composition and ratio of the lipids, particularly the Cers, determine the phase and nanostructure of the SC, and thus there has been proposed many different models to describe the lipid matrix observed from the experiments.^6^ Although the SC lipid organization is highly heterogenous, the coexistence of long- and short-repeat distance structures has been proposed.^15^ The short periodicity phase (SPP) and the long periodicity phase (LPP) have the repeat distances of approximately 6 nm and 13 nm, respectively. While the SPP is characterized as a lipid bilayer, the LPP is a more complex multilayer. A sandwich model, where two bilayers surround a fluid interior, was proposed for the LPP structure.^16^ 30-Linoyloxytriacontanoic acid-[(2S,3R)-1,3- dihydroxyoctadec-4-enyl]-amide (CerEOS), a Cer subclass containing a long fatty acid chain with linoleic acid, is known to be essential for the formation of the LPP structure,^15^ and a high CerEOS concentration is known to induce the fluidity of the interior.^17^ Moreover, the ^2^H-NMR analysis revealed that CerEOS can induce the formation of liquid domains in the SC matrix.^13^

A new class of epidermal Cers containing long acyl chains, called ‘1-*O*-acylceramides (CerEN)’, was relatively recently identified from human SC.^18^ It contains long acyl chains in both the *N-* and 1-*O*-positions, and thus more polar functional groups in the lipid head compared to other Cers. The physicochemical properties of CerENs indicate that they may play an important role in the organization of the SC lipid matrix and/or water barrier homeostasis, despite relatively low abundance (about 5 % of all esterified Cers or 2.3 % of all Cers).^18^ However, their precise contribution to the formation of the lipid matrix and the skin barrier remained to be unveiled.

Molecular dynamics (MD) simulation is a powerful tool to investigate the structure of the SC and interactions among lipid components at the atomistic level. To date, extensive studies have been performed to investigate the structures and properties of the SC^17,19–23^ including united-atom models^24–27^ and coarse-grained models.^28–31^ The SPP bilayers have been more extensively studied than the LPP multilayers due to their simpler structure. Although all-atom simulations of the LPP are limited, a recent study successfully implemented all-atom MD simulations to study molecular structures of the LPP in the SC with the sandwich model,^17^ providing insight into the formation of the LPP structure and behavior of the lipids.

In this study, we investigate the role of 1-*O*-acylceramide NP (CerENP) on the structure and permeability of the lipid matrix using the sandwich model (bilayer-slab-bilayer) of the LPP by all-atom MD simulations. We show that the splayed CerENP causes a significant change in the lipid matrix: disordering and lateral packing. Moreover, umbrella sampling simulations are performed on the LPP models to estimate the free energy and permeability to ethanol. Our study suggests that the CerENP with a long acyl chain plays roles in the organization of the lipid matrix and water barrier homeostasis.

## 2. EXPERIMENTAL SECTION

### System Setup

Two LPP models were constructed to investigate the lipid matrix (referred to as ‘Model A’ and ‘Model B’, respectively). The lipid compositions of these models were based on the previous report by reference 17, which shows a good agreement with an experimental model.^32^ These models may differ from a realistic model of the SC, because it is an isolated model based on the experimental lipid model.^32^ Despite, our results provide an insight into how a new subclass of ceramide effects on the structure and property of the SC. Both of the LPP models contain CerEOS, *N*-lignoceroylsphingosine (CerNS), *N*-oleoylphytosphingosine (CerNP), Chol, and lignoceric acid, while CerENP is included only into Model B (**Figure 1**). Lignoceric acid was used as a representative FFA. The exact compositions are provided in **Table 1**. Model A has the equimolar Cer:Chol:FFA ratio with a Cer mixture with the CerEOS:CerNS:CerNP ratio of 47:41:12. A higher concentration of CerEOS was used than the reported ratio of CerEOS to total Cers, ~5 mol% (or the ratio of overall ω-hydroxy Cers to total Cers, ~10 mol%)^33^ because ~30 % is the minimum concentration of CerEOS to exclusively form the LPP.^34^ Unlike the native SC that is in the coexistence of the LPP and SPP (and other multiple domains), the LPP alone structure used in this study might have different lipid concentrations, and thus the results should be interpreted with this uncertainty in mind. Model B contains the same number of lipids with additional CerENP (10 mol% of CerEOS), which slightly changes the lipid ratio; Cer:Chol:FFA ratio of 36:32:32, and CerEOS:CerNS:CerNP:CerENP ratio of 45:40:11:4, respectively. Although a relatively low abundance of 1-*O*-acylceramides (~5 % of all esterified Cers) has been reported for the mice epidermis,^18^ its concentration in human SC is unknown. Hence, the precise local concentration of CerENP, particularly in the LPP structure, is unclear. Here, we used 10 mol% of CerEOS by considering the high heterogeneity of the SC. CerENP was placed at the interface between the inner bilayer and the slab layer due to its splayed structure. For both of the models, five water molecules per lipid were placed, and no ions were included. Most water molecules were placed outside of the lipid matrix as an isolated model, and the water-to-lipid ratio is approximately 0.7, where it becomes about 2 if the first shell of water outside the bilayers is considered.^12,35^ CHARMM-GUI *Membrane Builder*^36,37^ was used to construct the models.

**Figure 1.**
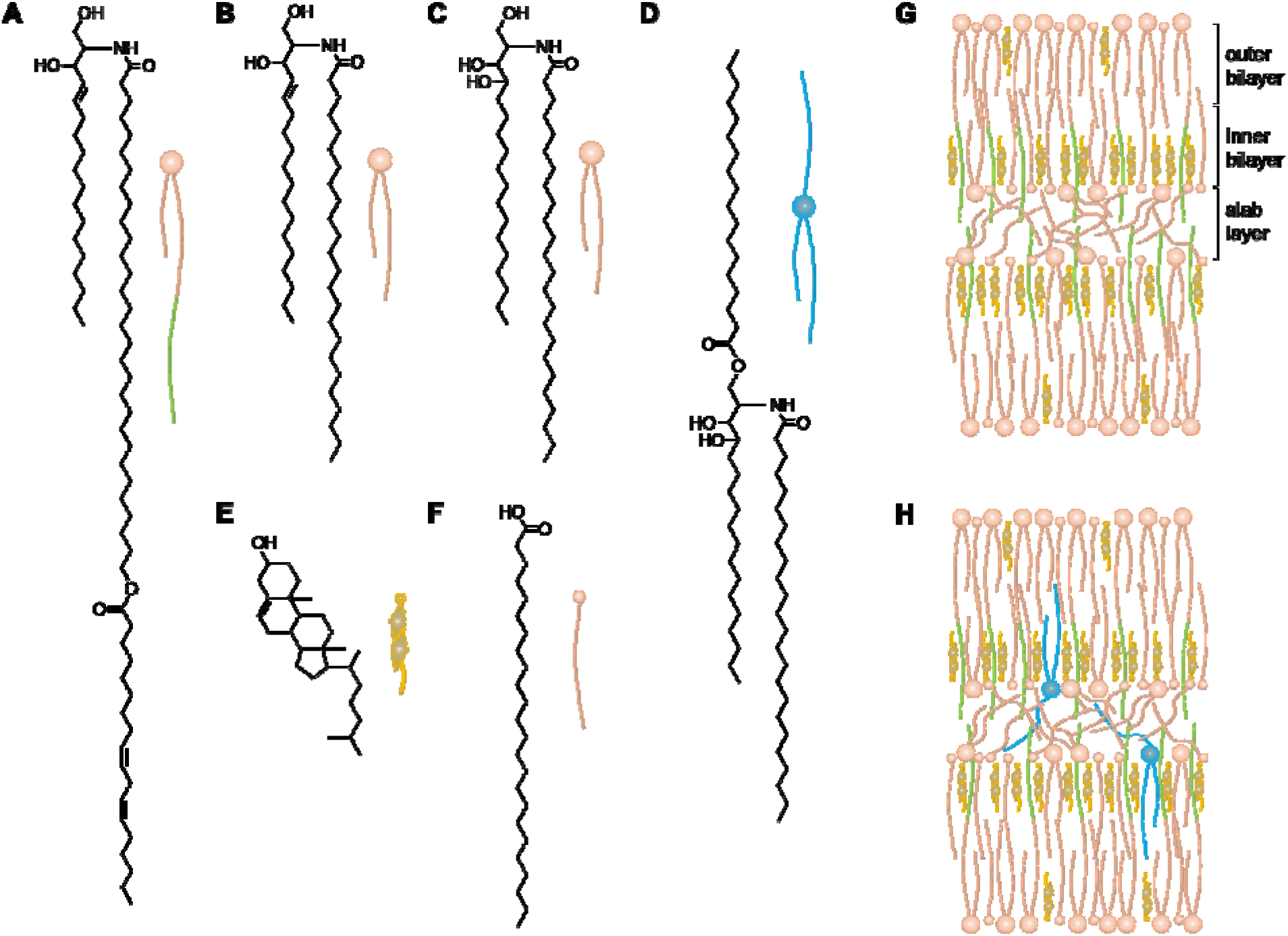
Chemical structures and schematic illustrations of (A) CerEOS, (B) CerNS, (C) CerNP, (D) CerENP, (E) Chol, (F) FFA, and lipid matrix (G) Model A and (H) Model B, respectively.

**Table 1.** Number of lipids per leaflet in Model A and Model B.

| Model<sup>a</sup> | Species | Outer bilayer | Inner bilayer | Slab layer |
| --- | --- | --- | --- | --- |
| A | CerNS | 15 | 20 | 36 |
|  | CerNP | 5 | 5 | 10 |
|  | CerEOS | 60 | 0 | 0 |
|  | Chol | 20 | 100 | 0 |
|  | FFA | 40 | 50 | 60 |
| B | CerNS | 15 | 20 | 36 |
|  | CerNP | 5 | 5 | 10 |
|  | CerEOS | 60 | 0 | 0 |
|  | CerENP | 0 | 6 | 0 |
|  | Chol | 20 | 100 | 0 |
|  | FFA | 40 | 50 | 60 |

<sup>a</sup>5 waters per lipid were included in each system, resulting in 3680 and 3740 waters in Model A and Model B, respectively.

### MD Simulations

All simulations were performed using GROMACS^38^ with the CHARMM36 lipid force field^39^ and the TIP3P water model.^40^ Steepest-descent energy minimization was performed first to relax the constructed system, which was further equilibrated by performing NVT (constant number of particles, volume, and 305 K temperature) and subsequent NPT (constant number of particles, 1 bar pressure, and 305 K temperature) simulations with positional restraints on the heavy atoms for 1 ns and 10 ns, respectively. The positional restraints were placed on the heavy atoms with a force constant of 9.6 kcal·mol^-1^·A□^-2^ during the NVT simulation. For the NPT simulation, the force constants were placed only on the lipid head groups to relax the lipid molecules, where the force constants were gradually reduced to 0 kcal·mol^-1^·A□^-2^ during the 10 ns NPT simulation. The time step was 2 fs, coordinates were saved every 10 ps, and product runs were carried out for ~900 ns for each model.

Umbrella sampling simulations were performed to investigate the permeability to ethanol. After the MD simulation, ethanol and additional water were added to the both models, which were followed by 10 ns MD simulations to relax the added ethanol and water molecules. To generate initial frames, we performed short restrained simulations, where an ethanol molecule was pulled along the *z* axis from the outside of the bilayer towards the center (the slab layer) with a force constant of 9.6 kcal·mol^-1^·A□^-2^. Trajectories of the retrained MD simulations are provided in Supporting Information (movies 1 and 2) to help understand how a molecule penetrates the lipid matrix. Note that these trajectories are not the result of umbrella sampling simulation. The window spacing was 1 A□ which produced 70 and 80 windows for Model A and Model B, respectively. For each window, 5 ns restrained simulations were performed with a force constant of 9.6 kcal·mol^-1^·A□^-2^. Five ethanol molecules were added to the LPP systems, and three independent umbrella sampling simulations were performed for three different ethanol molecules out of five. For each simulation, non-targeted four ethanol molecules were held in equidistant planes to prevent interactions. Potentials of mean force (PMFs) and diffusion profiles were calculated using the weighted histogram analysis method (WHAM).^41^

Electron density profiles (EDPs) were calculated using the SIMtoEXP^42^, and VMD^43^ program used for analysis and visualization.

## RESULTS AND DISCUSSION

### System Equilibration and Surface Area

We performed MD simulations to investigate the effect of CerENP on the lipid matrix structure. The surface area (SA), defined as the area of the simulation box (X·Y), as a function of time is useful to measure the system equilibration, and the calculated SA shows that the both model systems reached equilibration within the simulation time (**Figure 2A**). Each replicate of the both models also exhibited the similar result (Figure S1). The equilibrated total SA values were 58.8 nm^2^ and 54.7 nm^2^ for Model A and Model B, respectively. Interestingly, the SA value of Model B was significantly lower than that of Model A by ~7.5 % despite the greater number of lipids in Model B due to the additional CerENP. Also, the SAs per lipid, obtained by dividing the total surface area by the number of lipids in the outer bilayer leaflet, were 43.2 A□^2^ and 39.7 A□^2^ for Model A and Model B, respectively, which agree with previous studies.^17,20,21,44^ The change in the SA also affected the thickness of the lipid matrix: the thickness of the lipid matrix in Model B (13.3 nm) was larger than that in Model A (12.2 nm), which is also in agreement with the reported repeat distance of the LPP, ~13 nm. The SA difference between Model A and Model B indicates that the incorporated CerENP plays a significant role in the organization of the lipid matrix structure.

**Figure 2.**
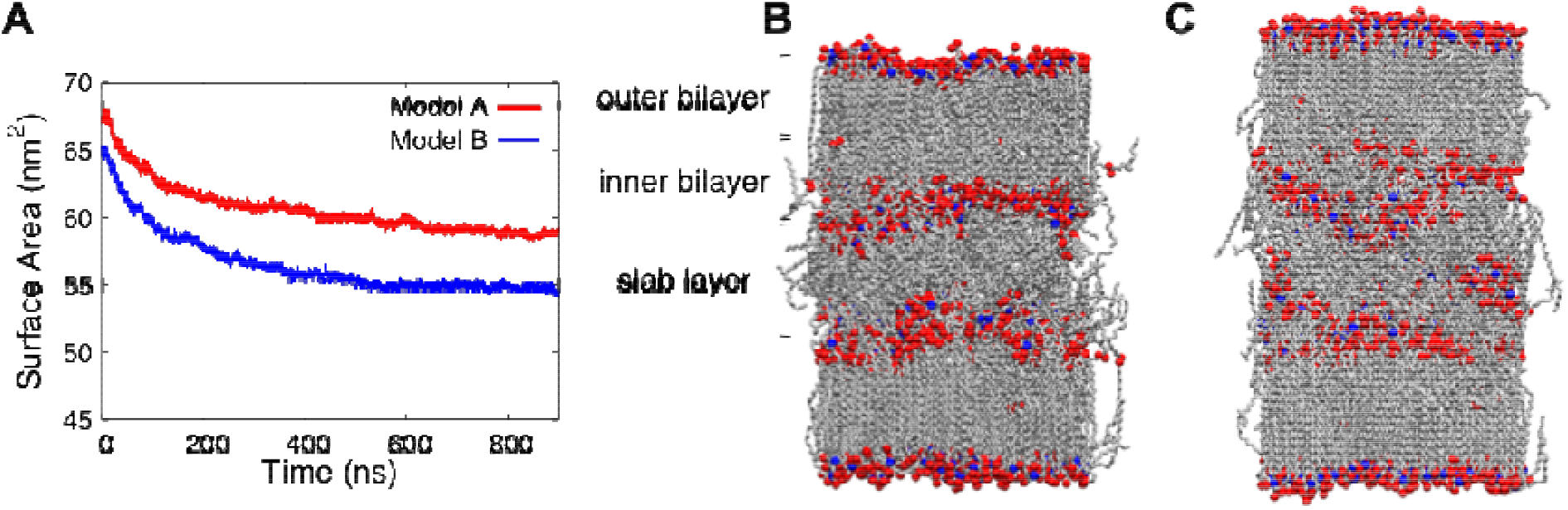
(A) The calculated total surface (SA) as a function of time for the both models. Snapshot structures of Model A (B) and Model B (C) at the end of simulation. Gray, blue, and red represent carbon, nitrogen, and oxygen atoms, respectively. Hydrogen atoms and water molecules are omitted for clarity.

The LPP model is composed of two bilayers surrounding a fluid interior slab layer with polar lipid headgroups in the both directions. Similar to the previous report,^10^ the simulation results indicate that the bilayer leaflets are relatively rigid, while the slab layer has a fluid-like structure in the both model systems (**Figure 2B**). However, the slab layer in Model B exhibited a relatively uneven interface with the bilayer leaflets, where the estimated standard deviations of the *z*-component for the head groups of lipids (tertiary carbons in Cer and carboxyl carbons in FFA) at the interface were 5.6 Å and 7.6 Å for Model A and Model B, respectively, indicating that the incorporation of CerENPs leads to the change in the lipid matrix structure

The calculated EDPs for the polar groups of lipids in Model A and Model B exhibit four peaks corresponding to interfacial regions (Figure S2A), which qualitatively agree with the previous experimental data by Groen et al.^45^ Note that the overall shape of the profiles is somewhat different from the experimental results because only polar groups were considered to clearly show the interfaces and locations of lipids. The head group of CerEOS is largely responsible for the outer peaks, while its ester group contributes to the inner peaks (Figures S2B and S2C). The inner peaks in Model B are slightly broadened as aforementioned, which is attributed to the shift of lipid locations, particularly CerNS, CerNP, and FFA. The distribution of each lipid species in the lipid matrix is further discussed in the following section.

### Molecular Distribution of the Lipid Matrix

The distribution of lipid molecules in each model at the end of simulation is presented in **Figure 3**. We also quantified the number of each lipid species in inner, outer, and slab layers before and after the simulation (Table S1), showing that a small number of Cers diffused across the layers, while many FFA and Chol molecules diffused across the interface, particularly from the inner bilayer to the slab layer.

**Figure 3.**
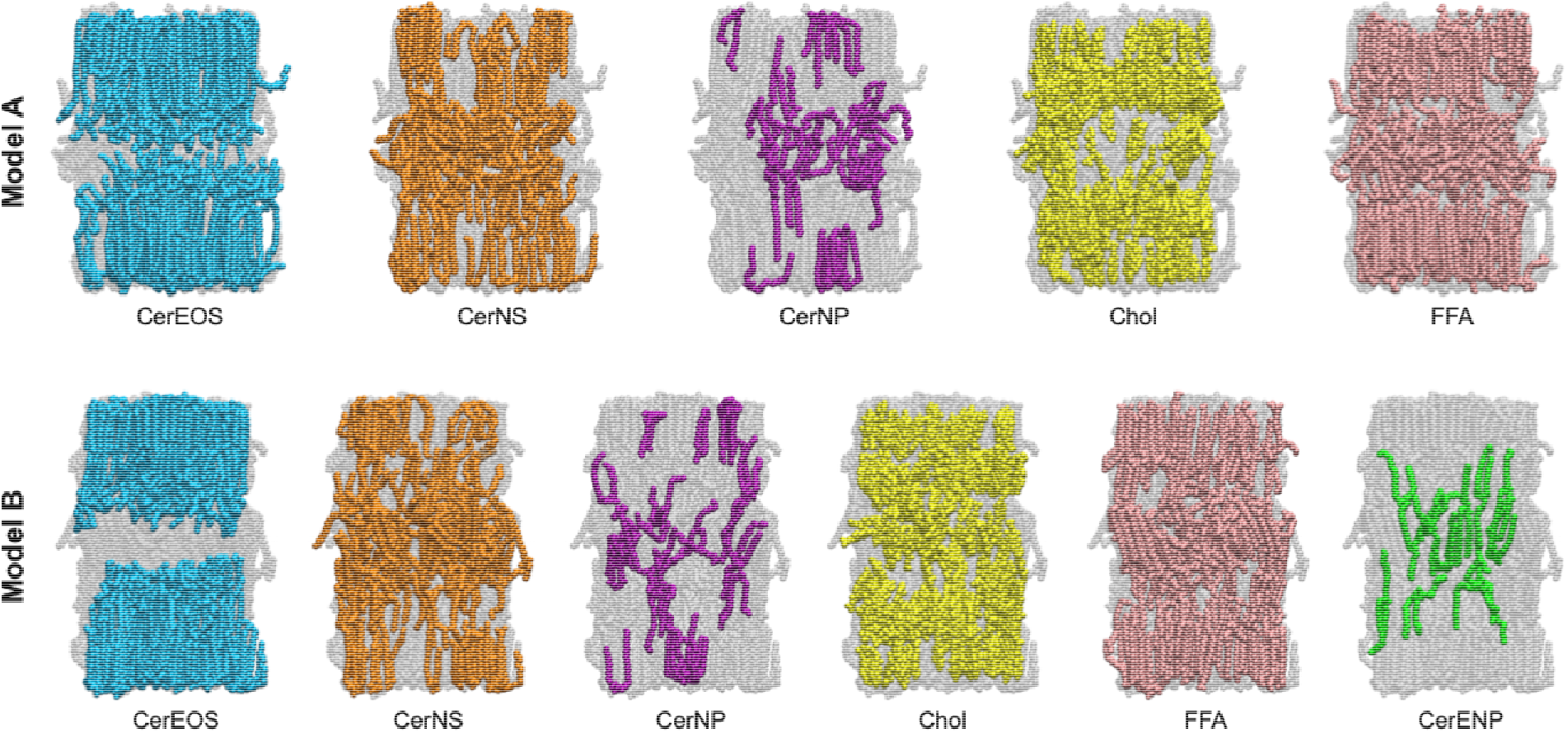
Distribution of each lipid component in Model A (upper) and Model B (lower), respectively, at the end of simulation.

In the LPP model proposed by Bouwstra et al.,^15^ CerEOS is one of the most important components in the lipid matrix formation as it occupies the outer bilayer leaflet and penetrates through the inner bilayer leaflet. It was also reported that a high CerEOS concentration (~8 mol%) induces fluidicity in the interior, while the interior is highly ordered at a lower concentration (~3 mol%).^10^ CerEOS has two double bonds in its linoleate tail of the fatty acid chain (Figure 1) and penetrates into the slab layer to bring a disorder in the interior. In the both models, the CerEOS molecules are ordered in the bilayer, whereas their linoleate tails of the fatty acid chain in the slab layer are disordered, contributing to the fluidity of the interior. The estimated order parameters (S_CD_) of CerEOS show the higher values for the sphingosine and fatty chains (S_CD_ > 0.4), whereas significantly lower values for the linoleate tails of the fatty acid chain in the slab layer are disordered (S_CD_ < 0.2) (Figure S3). Model B exhibits a relatively compact and dense distribution owing to the smaller SA compared to that of Model A. Also, as Model B has a thicker lipid matrix, a significant gap exists between the CerEOS tail ends from the upper bilayer and the lower bilayer, compared to those of Model A, where they are positioned close to each other in the slab layer.

CerNS and CerNP (24-carbon Cer) can take two different conformations in the lipid matrix: hairpin (lipid chains point in same direction) and extended (chains point in opposite directions) conformations (Figure S4). Although we started the simulations with the hairpin conformation for all of the Cer lipids, a significant number of the lipids, about 13 % and 18 % of total 24-carbon Cers for Model A and Model B, respectively, were changed to the extended conformation during the simulations (Figure S4). The extended conformation of Cers was reported from both experiments and simulations,^14,17,46^ indicating that it may contribute to the heterogeneity of the lipid matrix. Also, the snapshot structures exhibit different conformations depending on the position, rigid and disordered structures in the bilayer and in the slab layer, respectively (Figure 3).

The Chol and FFA molecules are abundant and located over the lipid matrix (Figure 3). Although the Chol molecules were first placed mainly in the inner bilayer, we observed that ~15 % (Model A) and 14 % (Model B) of total Chol molecules migrated across the interface between the bilayer and the slab layer during the simulations, while a few Cers migrated across the interface (Table S1). FFA, similar to the CerNS and CerNP, shows rigid structures in the bilayer and disordered structures in the slab layer, respectively. In fact, FFA is considered to diffuse through the lipid matrix due to its simple single-chain structure. For this reason, it was suggested that FFA is regulated by transport proteins associated with lipid rafts.^47^

### Hydrogen Bond Interactions

The lipids form hydrogen bonds (H-bonds) between the lipids due to the polar head groups, contributing to the formation of the lipid matrix. In particular, sphingolipids that contain numerous H-bond groups capable of acting as both donors and acceptors. **Table 2** provides the average number of H-bonds per a lipid for each lipid species. The result indicates that the lipids in Model B form more H-bonds than Model A up to 0.1 per a lipid, which is attributed to the smaller SA or the lateral chain packing in the lipid matrix. However, it has been reported that the increased H-bonds do not always lead to a denser packing due to the increase of SA in head groups,^48^ so the relationship between the number of H-bonds and the packing of the lipid matrix should be carefully considered. Cer species show a relatively large number of H-bonds than Chol or FFA due to their polar groups with multiple H-bond donor and acceptor sites. Interestingly, CerENP exhibits the greatest number of H-bonds compared to other Cers. Indeed, these values are roughly proportional to the number of H-bond donors/acceptors: CerEOS, CerNS, CerNP, and CerENP contain 4, 4, 5, and 6 H-bond donors/acceptors in the head group, respectively. Thus, the splayed CerENP can form multiple H-bonds with neighboring lipids (**Figure 4**). The results indicate that CerENP can contribute to the organization of the lipid matrix despite its relatively low abundance in the skin.

**Figure 4.**
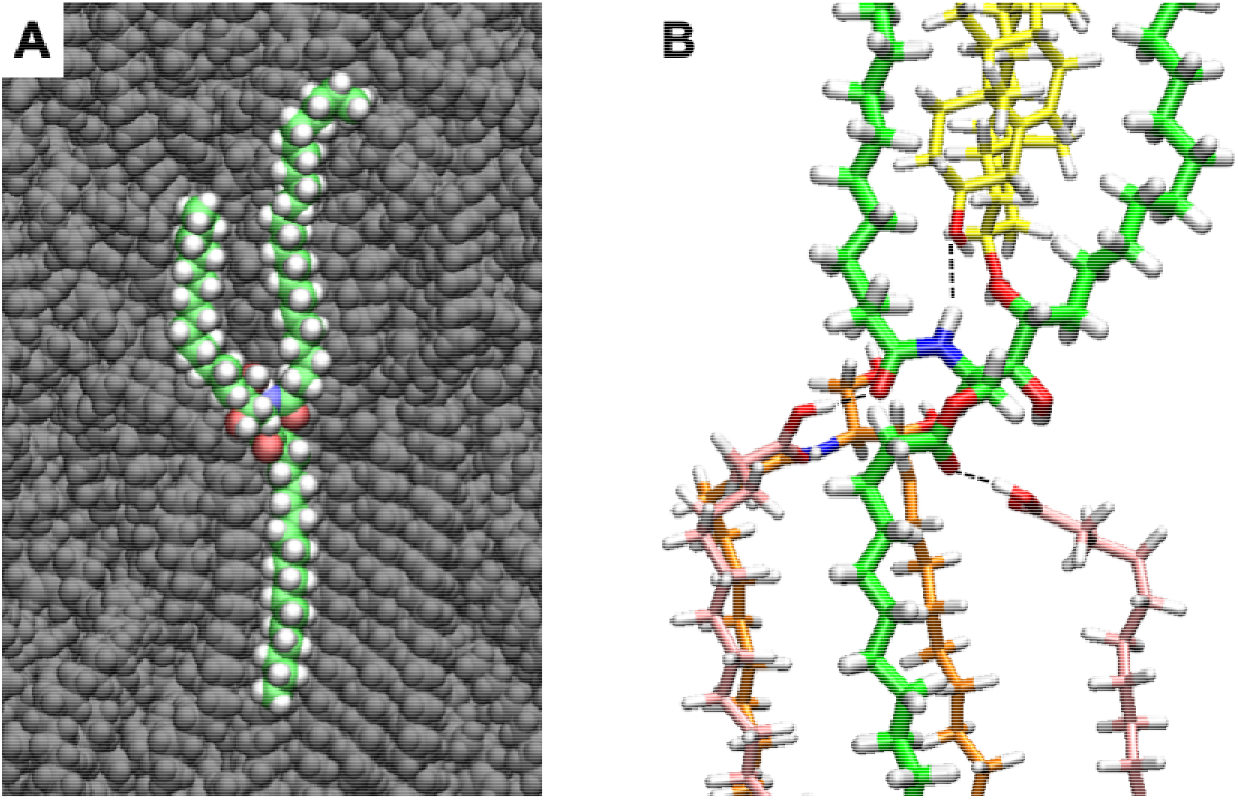
(A) Snapshot of CerENP in the lipid matrix. (B) The polar interactions of CerENP with other lipids, where carbon atoms in CerENP, CerNS, Chol, and FFA are shown in green, orange, yellow, and pink, respectively. Red, blue, white, and black dotted lines represent oxygen, nitrogen, hydrogen, and hydrogen bonds, respectively.

**Table 2.**
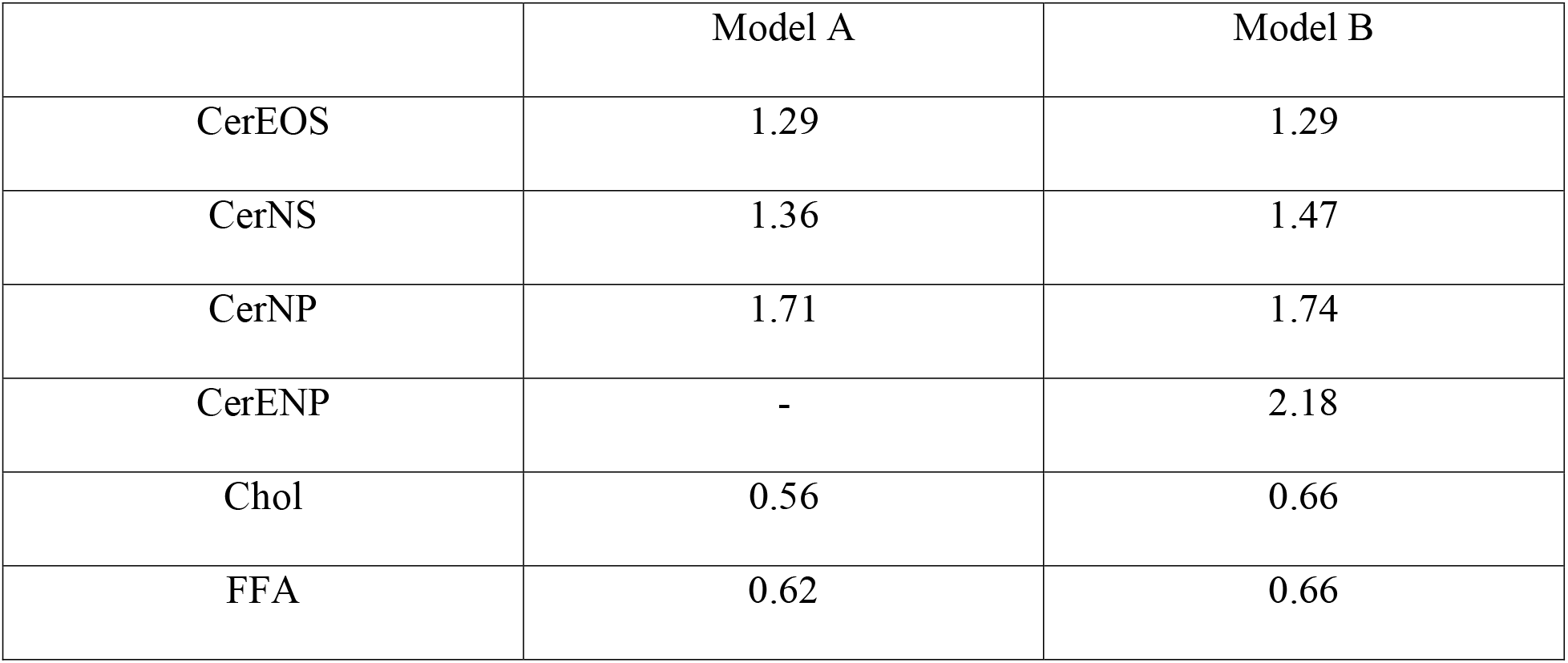
Average number of H-bonds per one lipid between the lipids for each lipid species, where intramolecular H-bonds were excluded.

We further estimated the average number of H-bonds for each pair of the lipid species in the structure to investigate specific interactions by lipid species (Tables S2 and S3). The number of H-bonds by lipid species is roughly proportional to the number of the lipids. As the head group of CerEOS is located in the outer bilayers, it forms H-bonds with mostly FFA and other CerEOS molecules. In comparison, CerNS is positioned mainly in the inner bilayers and the slab layer, and thus it forms H-bonds with mainly Chol and FFA species. CerNP and CerENP form a small number of H-bonds with other lipids due to their relative low abundance, and Chol and FFA form H-bonds roughly proportional to the number of lipid species because they are distributed over the whole lipid matrix.

### Permeability to Ethanol

To investigate the effect of CerENP on the diffusion of a molecule penetrating the lipid matrix, we performed umbrella sampling simulations for the both Model A and Model B, where ethanol was used as a penetrating molecule, a common chemical enhancer used in topical formulations to increase the penetration.^49^ The PMFs of ethanol as a function of position along the lipid matrix (the *z*-axis) were estimated for the both models (**Figure 5**). Model A exhibits a large peak at around 4.1 nm with 7.8 ± 1.7 kcal mol^-1^, and the values are quite similar to the previous report.^10^ In comparison, Model B shows a large peak at around 4.5 nm with 9.2 ± 2.1 kcal mol^-1^, which is attributed to the thicker and denser lipid matrix than Model A. The higher energy for the penetration of the ethanol molecule might result from the hydrophobic hydrocarbons of the lipids. The results indicate that CerENP induces lateral packing in the lipid matrix, which may increase the barrier function against the penetration of ethanol molecules.

**Figure 5.**
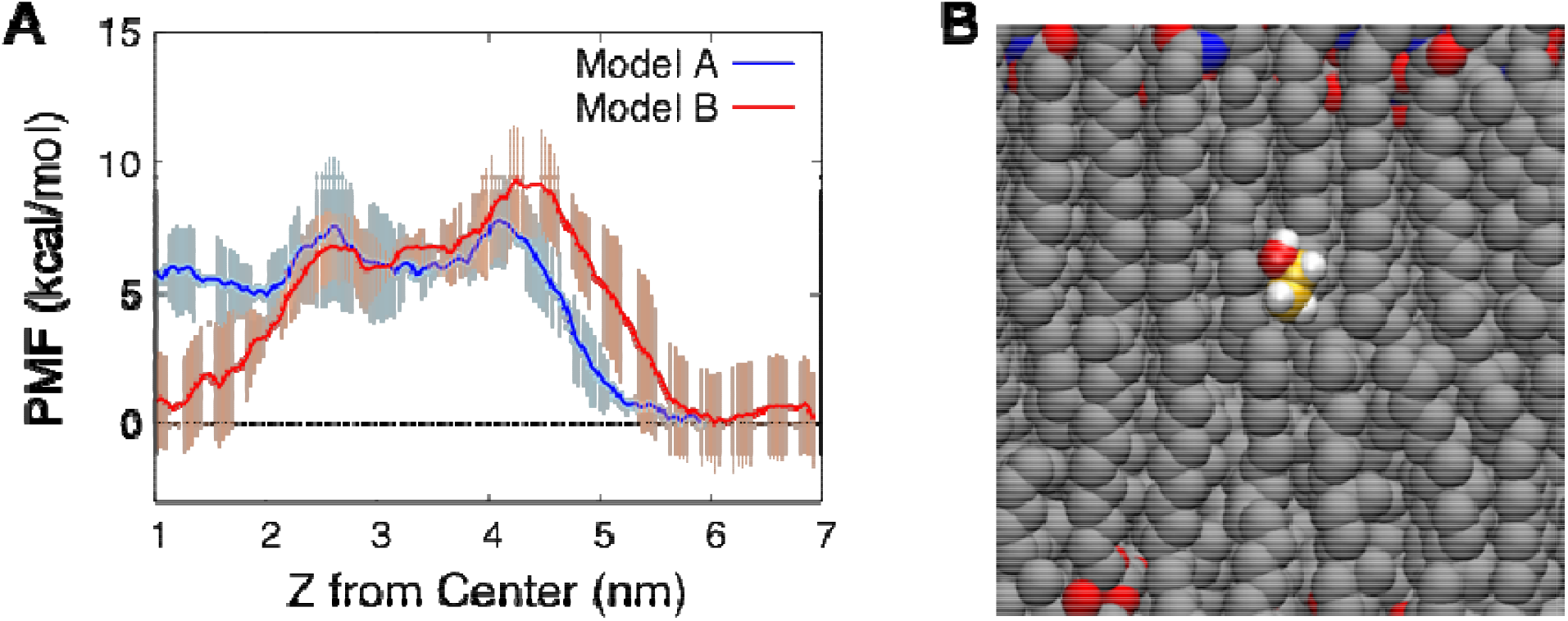
(A) PMFs for ethanol permeation through the lipid bilayer in Model A and Model B, where standard errors (thin lines) were estimated from three different calculations for each model. (B) A snapshot structure of ethanol in the lipid matrix, where the carbon atoms of ethanol and lipids are colored in gold and gray, respectively. Oxygen, nitrogen, and hydrogen atoms are red, blue, and white, respectively, where hydrogens in the lipids are omitted for clarity.

## CONCLUSIONS

We carried out microsecond MD simulations of the LPP model to investigate the role of CerENP on the structural organization and permeability of the SC lipid matrix. The lipid matrix with the bilayer-slab-bilayer structure was constructed using CerEOS, CerNS, CerAP, Chol, and LA. The addition of CerENP to the SC lipid matrix induced disordering at the bilayer/slab interface and lateral lipid packing. CerENP possess a number of H-bond donors/acceptors in the structure, leading to multiple H-bonds with neighboring lipids. Moreover, the presence of CerENP increases the barrier function of the lipid matrix against ethanol penetration, which implies that the structural modification of the lipid matrix induced by a small number of CerENP can have a significant impact on the skin barrier function. This study demonstrated that MD simulations can provide valuable insight into the biological role of a new subclass of Cer in the SC and allow us to better understand how various lipid structures contribute to the skin organization.

## ASSOCIATED CONTENT

### Supporting Information

The Supporting Information is available free of charge. Surface area plot for the replica calculations, the electron density profiles, the order parameters, the extended conformations of CerNS and CerNP, and the average number of H-bonds by lipid species (PDF). Trajectories of the restrained MD simulations for ethanol penetration (movies 1 and 2).

## AUTHOR INFORMATION

### Notes

The authors declare no competing interests.

## ACKNOWLEDGMENT

This research was supported by a grant from the Korea Health Technology R&D Project through the Korea Health Industry Development Institute (KHIDI), funded by the Ministry of Health & Welfare, Republic of Korea (grant number: HP20C0018). Computations were supported by the National Institute of Supercomputing and Network/Korea Institute of Science and Technology Information with supercomputing resources including technical support (KSC-2018-S1-0004).

