## Supplementary information for "The Role of 1-*O*-Acylceramide NP in Structural Organization and Permeability of the Stratum Corneum Lipid Matrix"

**This document includes:**

Figures S1 to S3

Tables S1 to S3

**Other supplementary materials for this manuscript include the following:**

Movies S1 to S2

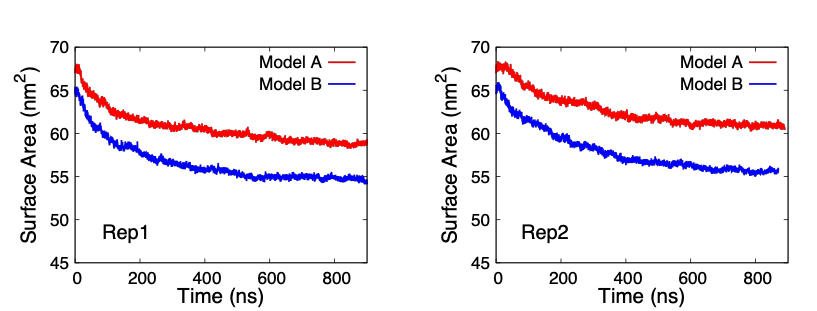

**Figure S1.** Comparison of total SA as a function of time for Models A and B. Each model has two replicates.

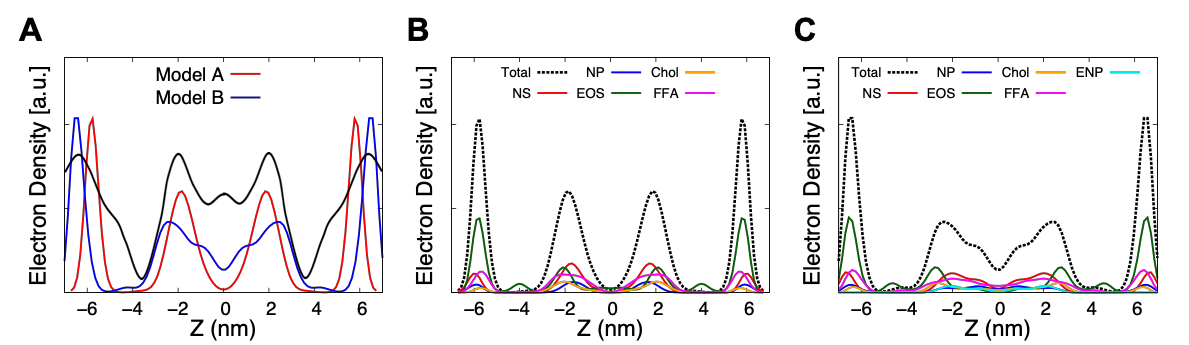

**Figure S2.** (a) The calculated EDPs for polar groups of total lipid in Model A (red) and Model B (blue), where the black line represent the experimental data reported by Groen et al.^1^ Polar EDPs for each lipid species in (B) Model A and (C) Model B, respectively. The center of the lipid matrix corresponds to the Z=0 nm.

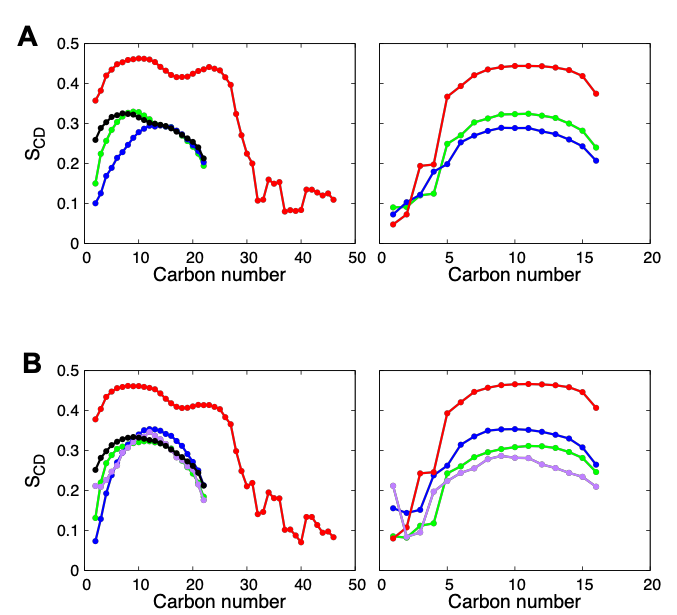

**Figure S3.** Order parameters (S_CD_) as a function of chain carbon number for fatty acid chain (left) and sphingosine chain (right) of Cer and FFA for (A) Model A and (B) Model B, respectively. Black, red, blue, green, and purple represent FFA, CerEOS, CerNP, CerNS, and CerENP, respectively.

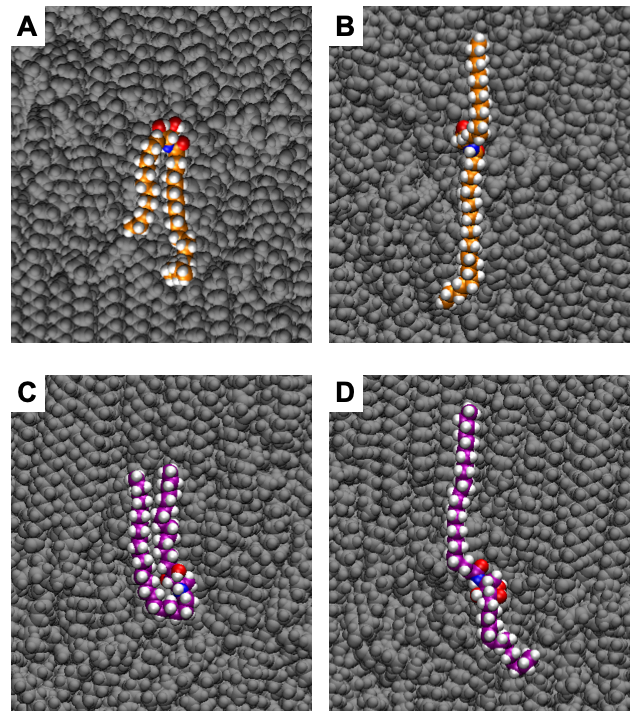

**Figure S4.** Snapshot structures of CerNS with (A) the hairpin and (B) the extended conformations, and CerNP with (C) the hairpin and (D) the extended conformations, respectively. Carbon atoms of CerNS and CerNP are shown with orange and purple, and red, blue, and white represent oxygen, nitrogen, and hydrogen, respectively.

**Table S1.** Number of lipids per leaflet in Model A and Model B after the 900 ns MD simulations (numbers in parentheses indicate the number of lipids in the initial structures).

|  | **Species** | **Outer bilayer** | **Inner bilayer** | **Slab layer** |
| --- | --- | --- | --- | --- |
| **Model A** | CerNS | 15 (15) | 18 (20) | 40 (36) |
|  | CerNP | 5 (5) | 4 (5) | 12 (10) |
|  | CerEOS | 60 (60) | 0 (0) | 0 (0) |
|  | Chol | 21 (20) | 82 (100) | 34 (0) |
|  | FFA | 40 (40) | 41 (50) | 78 (60) |
| **Model B** | CerNS | 15 (15) | 20.5 (20) | 35 (36) |
|  | CerNP | 5 (5) | 4 (5) | 12 (10) |
|  | CerEOS | 60 (60) | 0 (0) | 0 (0) |
|  | CerENP | 0 (0) | 5 (6) | 2 (0) |
|  | Chol | 27.5 (20) | 75.5 (100) | 34 (0) |
|  | FFA | 40 (40) | 42.5 (50) | 75 (60) |

**Table S2.** Average number of H-bonds by lipid species for Model A, where intramolecular H-bonds were excluded.

|  | **CerEOS** | **CerNS** | **CerNP** | **Chol** | **FFA** |
| --- | --- | --- | --- | --- | --- |
| **CerEOS** | 58.66 | 17.80 | 13.82 | 25.64 | 38.55 |
| **CerNS** | 17.80 | 16.95 | 10.67 | 50.46 | 48.26 |
| **CerNP** | 13.82 | 10.67 | 2.45 | 13.17 | 11.28 |
| **Chol** | 25.64 | 50.46 | 13.17 | 7.35 | 37.00 |
| **FFA** | 38.55 | 48.26 | 11.28 | 37.00 | 12.57 |
| **Total** | 154.47 | 144.14 | 51.39 | 133.62 | 147.66 |

**Table S3.** Average number of H-bonds by lipid species for Model B, where intramolecular H-bonds were excluded.

|  | **CerEOS** | **CerNS** | **CerNP** | **CerENP** | **Chol** | **FFA** |
| --- | --- | --- | --- | --- | --- | --- |
| **CerEOS** | 60.94 | 13.42 | 9.29 | 1.50 | 29.68 | 40.40 |
| **CerNS** | 13.42 | 21.06 | 8.90 | 9.89 | 54.96 | 47.29 |
| **CerNP** | 9.29 | 8.90 | 2.57 | 4.55 | 14.39 | 12.53 |
| **CerENP** | 1.50 | 9.89 | 4.55 | 0.00 | 5.99 | 4.28 |
| **Chol** | 29.68 | 54.96 | 14.39 | 5.99 | 9.10 | 45.25 |
| **FFA** | 40.40 | 47.29 | 12.53 | 4.28 | 45.25 | 8.80 |
| **Total** | 155.23 | 155.52 | 52.23 | 26.21 | 159.37 | 158.55 |
